# Decrease in *ACE2* mRNA expression in aged mouse lung

**DOI:** 10.1101/2020.04.02.021451

**Authors:** A. Sina Booeshaghi, Lior Pachter

## Abstract

Angiotensin-converting enzyme 2 (*ACE2*) has been identified as a critical receptor for severe acute respiratory syndrome coronavirus 2 (SARS-CoV-2). This has led to extensive speculation on the role of *ACE2* in disease severity, and in particular, whether variation in its expression can explain higher mortality in older individuals. We examine this question in mouse lung and show that 24-month old mice have significantly reduced *ACE2* mRNA expression relative to 3-month old mice. The differences appear to be localized to ciliated cells.

## Introduction

The severity of illness resulting from infection with SARS-CoV-2 virus ^1^ is strongly correlated with age ^2^. There has been much speculation about the mechanisms underlying this strong association, with one hypothesis being that age-related changes in *ACE2*, a crucial receptor used by SARS-CoV-2 to enter cells ^3^, affect disease susceptibility ^4^. Nevertheless, there has been little data on the age dependence of *ACE2* expression in humans, with the exception of GTEx data, which is limited in the age range to adults ^5^.

To better assess the relationship between age and *ACE2* expression, we examined single-cell RNA-seq data from eight 3-month old and seven 24-month old mouse lungs ^6^ in order to identify cell-type specific age-related differences, if any, in *ACE2* mRNA expression. Single-cell RNA-seq can determine whether bulk shifts in gene expression are due to changes in cell-type abundances, or changes in gene expression within specific cell types ^7^. We find that both cell-type abundance shifts, as well as *ACE2* mRNA changes in specific cell types, contribute to a decrease in *ACE2* mRNA with age. We corroborate our findings with bulk RNA-seq analysis and *ACE2* protein measurement in rat lungs ^8^.

## Results

To examine mRNA abundance of *ACE2* in young and old mice lungs, we began by uniformly processing ^9,10^ all RNA-seq from GSE12487 ^6^ (see Methods), followed by computation of common coordinates for the 3-month and 24-month old mice single-cell RNA-seq using scVI ^11^ (Figure 1a, Methods). Consistent with previous findings ^12^, we find that *ACE2* is not prevalent in the lung, with the exception of Goblet/Club and ciliated cells (Supp. Fig. 1). In cells where *ACE2* mRNA molecules are identified, the average copy number is 1.19. We, therefore, measured the expression level of *ACE2* by the proportion of cells with at least one copy (Methods).

**Figure 1:**
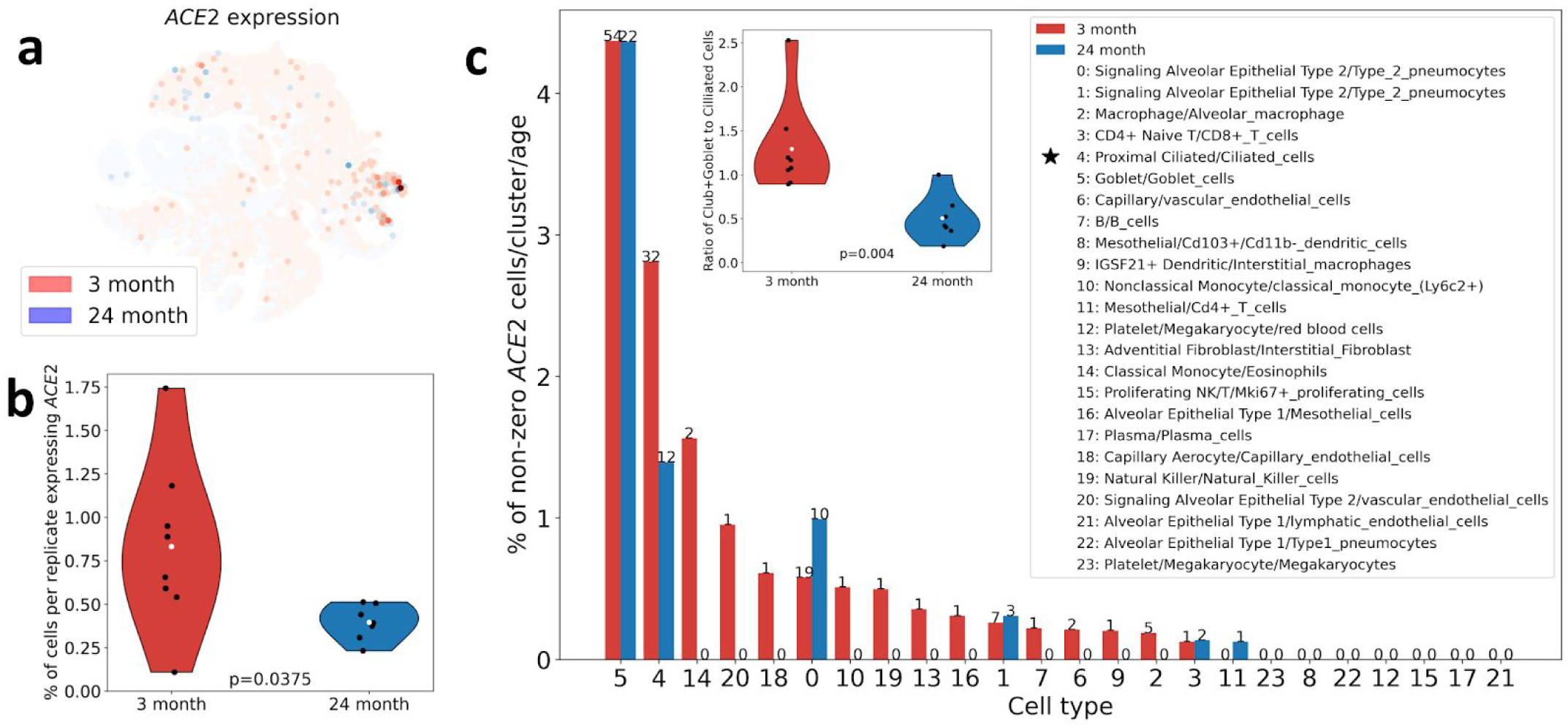
a) *ACE2* mRNA expression in a tSNE representation of scVI coordinates for 3-month and 24-month old mice. b) Proportion of cells displaying *ACE2* in 3-month old vs. 24-month old mice. c) *ACE2* mRNA expression by cell type in young vs. old mice (inset: ratio of Goblet/Club cells to ciliated cells in 3-month old vs. 24-month old mice).

In a comparison of the 3-month old and 24-month old mice, we found significantly fewer cells with *ACE2* expression in the 24-month old mice (2.1 fold reduction, p = 0.0375, Figure 1b). We corroborated this result with an analysis of three bulk RNA-seq samples from 3-month old mice and three bulk RNA-seq samples from 22-month old mice (1.5 fold reduction, p = 0.0163, Supp. Fig. 2). This reduction in *ACE2* mRNA expression is also seen in an analysis of FACS sorted alveolar macrophage bulk RNA-seq samples (2.7 fold reduction, p = 0.3076, Supp. Fig 3). These results are consistent with a previously observed dramatic reduction of *ACE2* protein between 3-month old and 24-month old rat lungs ^8^.

The decrease in *ACE2* mRNA expression with age in lungs is likely due to two underlying phenomena: a reduction of *ACE2* mRNA in ciliated cells, and a shift in ciliated cell abundance with age (Figure 1c, Supp. Fig 4). The latter result is statistically significant and corroborated in an analysis of FACS sorted cells ^6^.

SARS-CoV-2 uses the serine protease *TMPRSS2* for Spike protein priming, so we repeated our analyses for *TMPRSS2*. We found no significant changes in mRNA expression between 3-month and 24-month old mice (Supp. Figs. 5-7).

## Discussion

Our results show a decrease in *ACE2* mRNA expression between young and old mice and contradict recent claims by others ^13^. To understand the discrepancies in results, we examined differences in data or methodology that could lead to contradictory results (Supp. Table 1). We conclude that there is strong evidence in support of a decrease in *ACE2* mRNA between young and old mice, and that other currently available RNA-seq datasets do not contradict this finding. We note that results in rodents may not necessarily translate to humans.

## Methods

All methods and code to generate the results in this manuscript can be found in the python notebooks at https://github.com/pachterlab/BP_2020.

## Data availability

Raw FASTQ files can be obtained from <u>GSE124872</u>.

## Notes

https://github.com/pachterlab/BP_2020

