## Supplementary Figures for "Decrease in *ACE2* mRNA expression in aged mouse lung"

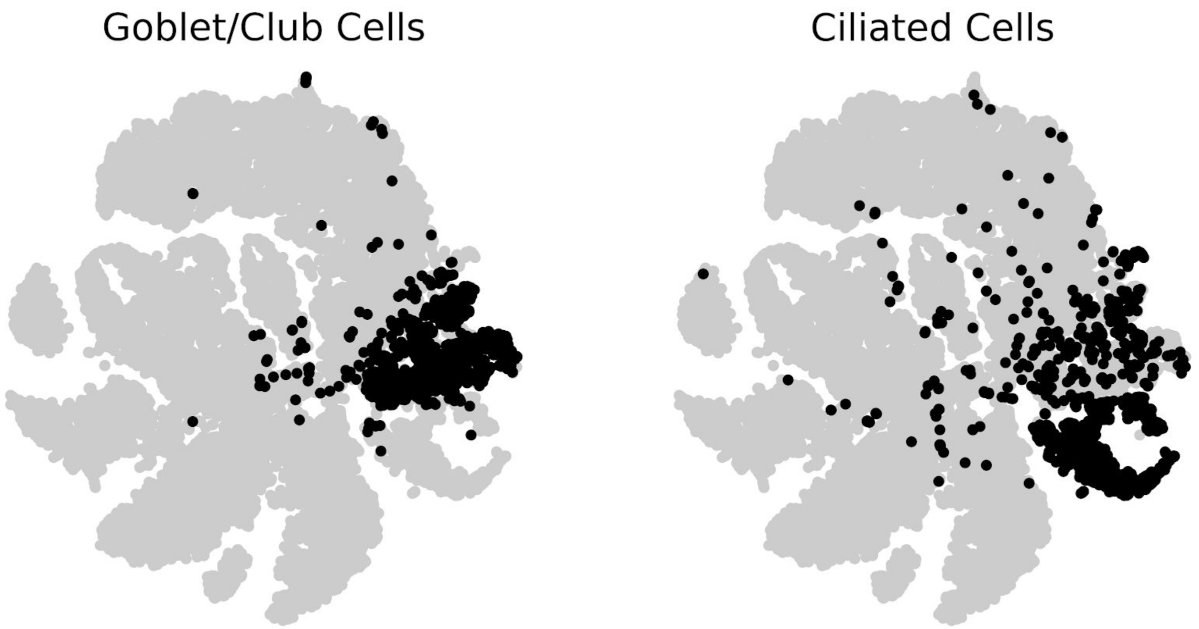


**Supplementary Figure 1:** t-distributed Stochastic Neighbor Embedding of 10 scVI components showing cell type in black.

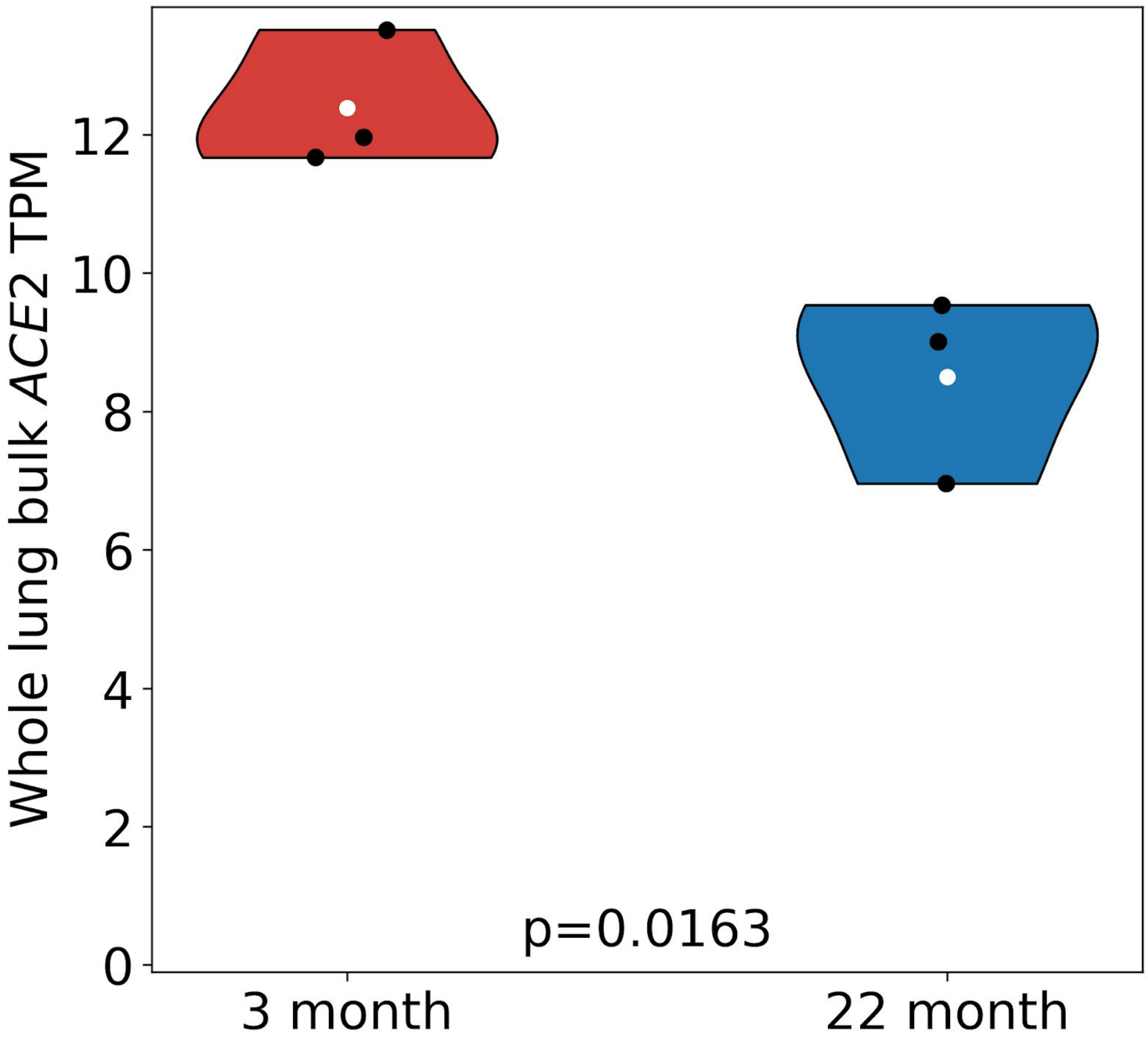


**Supplementary Figure 2:** Comparison of *ACE2* mRNA expression in 3-month old and 22-month old whole lung bulk RNA-seq.

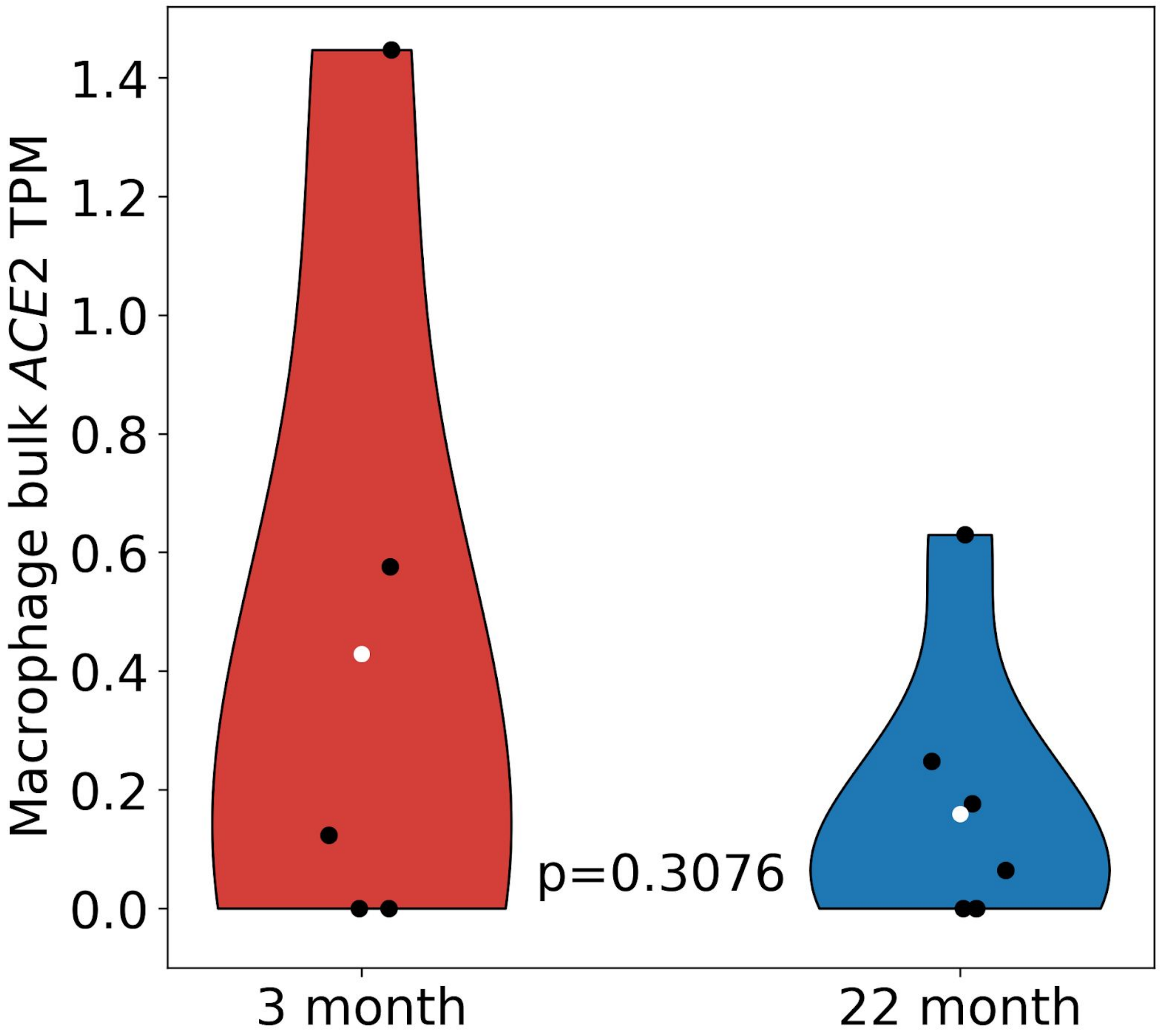


**Supplementary Figure 3:** Comparison of *ACE2* mRNA expression in 5 3-month old and 6 22-month old alveolar macrophage bulk RNA-seq.

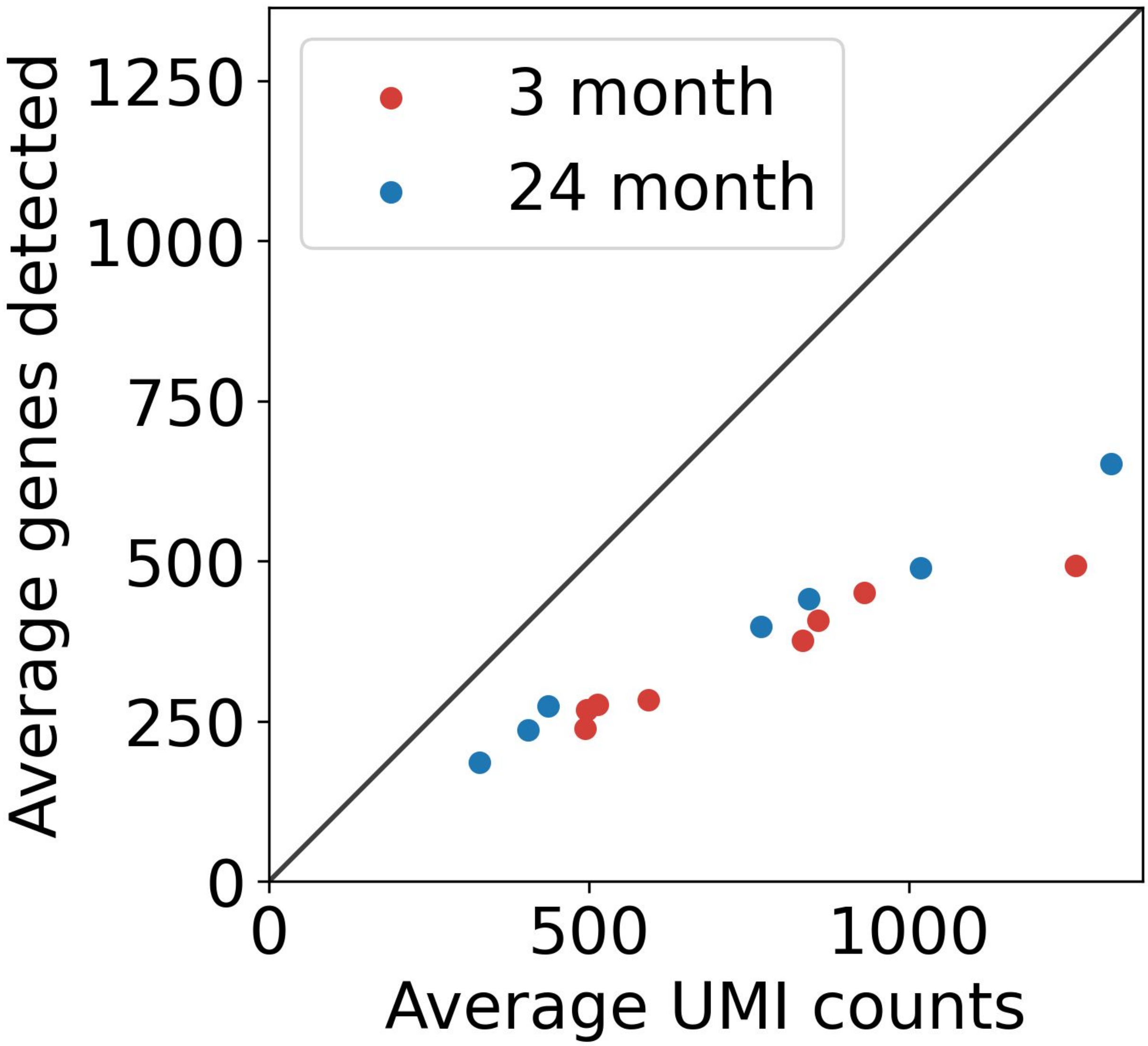


**Supplementary Figure 4:** Average number of UMIs and genes detected per cell colored by sample. There is no significant difference between 3-month old and 24-month old samples .

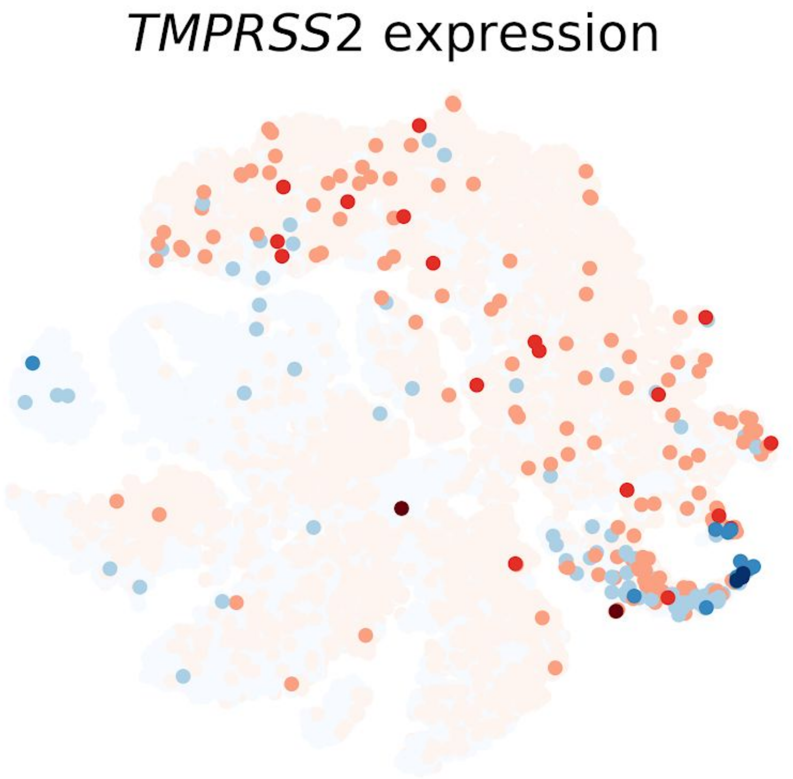


**Supplementary Figure 5:** t-distributed Stochastic Neighbor Embedding showing cells that express *TMPRSS2* (3-month old cells in red, 24-month old cells in blue).

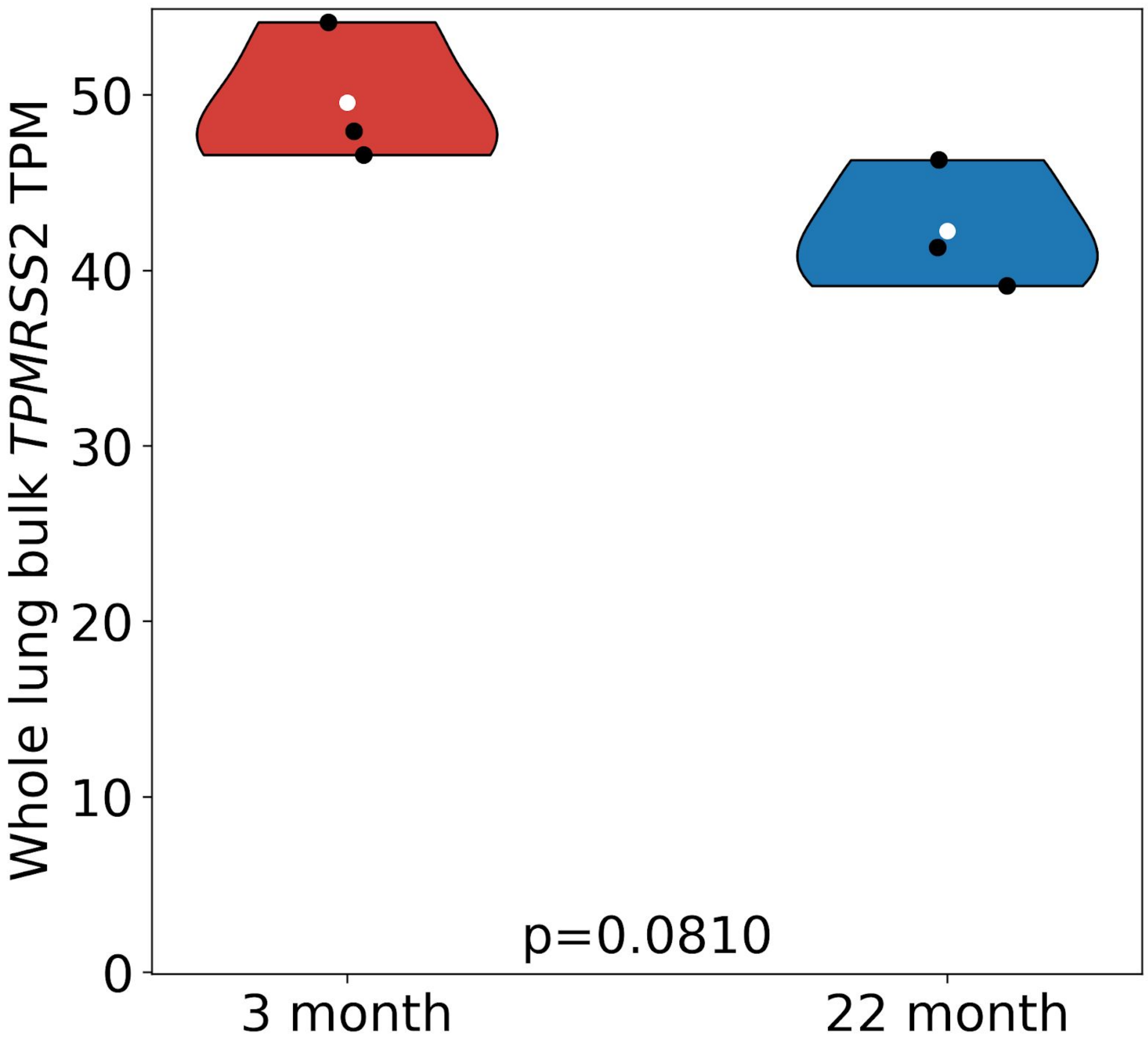


**Supplementary Figure 6:** Comparison of *TMPRSS2* mRNA expression in 3-month old and 22-month old whole lung bulk RNA-seq.

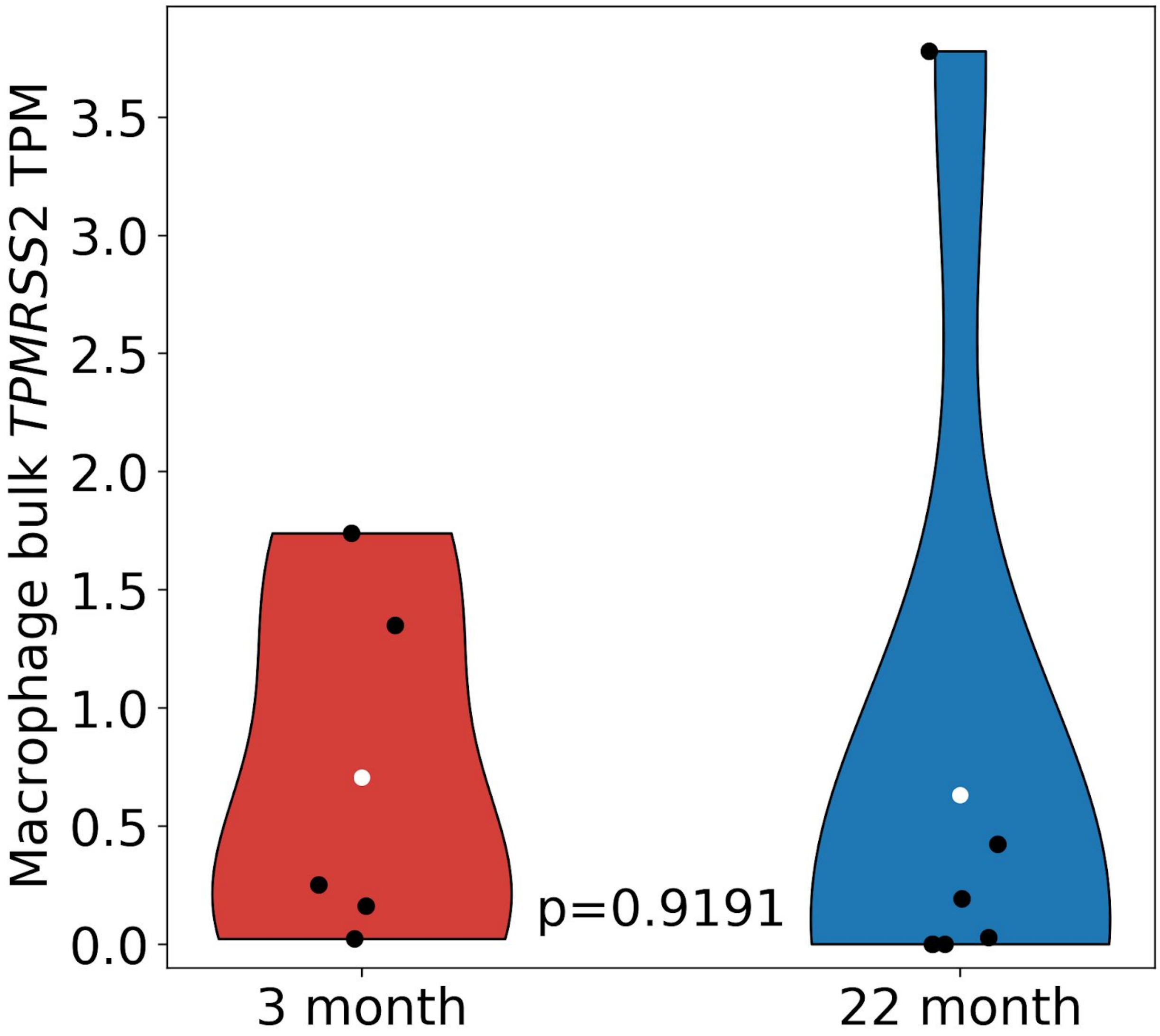


**Supplementary Figure 7:** Comparison of *TMPRSS2* mRNA expression in 5 3-month old and 6 22-month old alveolar macrophage bulk RNA-seq.

**Supplementary Table 1**: Claims that *ACE2* mRNA expression does not decrease with age.

| Dataset | Claim | Refutation |
|---|---|---|
| GTEx[1] | Chen *et al*[2]:"*ACE2* expression generally decreases with age significantly or insignificantly." <br><br> Smith and Sheltzer[3]: "*ACE2* expression [in GTEx] was equivalent between...young individuals (<29 years) and elderly individuals (>70 years)." | Multiple confounders that are difficult to correct for; RNA from cadavers may be of degraded quality; despite these issues evidence of slight decrease in *ACE2* expression with age. |
| Rat BodyMap[4] | Smith and Sheltzer[3]: "...young rats (6 weeks) and elderly rats (104 weeks) [displayed equivalent levels of *ACE2* expression in the lung]" | Single-end 50bp bulk total RNA was quantified with Cufflinks v2.0.2[5] (2012) which did not perform multiple rounds of EM for reads ambiguous between genes. *ACE2* is a member of a gene family. Furthermore, the Ribo-zero requires careful normalization. |
| Bulk RNA-seq from mice alveolar macrophages[6] | We find that 22-month samples have increased expression of *ACE2* on average (0.47 fold change young/old p=0.1198). (Methods) | Increased *ACE2* expression in 24-month lung in FACS sorted bulk Epithelium (Club/Ciliated) is partly explained by different proportions of club/goblet cells to ciliated cells in comparison with proportions from single-cell RNA-seq (Figure 1c). |
